# The psychometric validation of the Dutch version of the Rivermead Post-Concussion Symptoms Questionnaire (RPQ) after Traumatic Brain Injury (TBI)

**DOI:** 10.1101/502534

**Authors:** Anne Marie C Plass, Dominique Van Praag, Amra Covic, Anastasia Gorbunova, Ruben Real, Nicole von Steinbuechel, Dutch and Flemish CENTER TBI investigators.

## Abstract

**Background:** Traumatic brain injury (TBI) is one of the most common neurological conditions. It can have wide-ranging physical, cognitive and psychosocial effects. Most people recover within weeks to months after the injury, but a substantial proportion are at risk of developing lasting post-concussion symptoms. The Rivermead Post-Concussion Syndrome Questionnaire (RPQ) is a short validated 16-items self-report instrument to evaluate post-concussive symptoms. The aim of this study was to test psychometrics characteristics of the current Dutch translation of the RPQ.

**Methods:** To determine the psychometric characteristics of the Dutch RPQ, 472 consecutive patients six months after they presented with a traumatic brain injury in seven medical centers in the Netherlands *(N*=397), and in two in Belgium (Flanders) (*N*=75) took part in the study which is part of the large prospective longitudinal observational CENTER-TBI-EU-study. Psychometric properties at six months post TBI, were assessed using exploratory and confirmatory factor analyses. Sensitivity was analyzed by comparing RPQ scores of patients after mild vs. moderate and severe TBI.

**Findings:** The Dutch version of RPQ proved good, showing excellent psychometric characteristics: high internal consistency (Cronbach’s α .93), and good construct validity, being sensitive to self-reported recovery status at six months post TBI. Moreover, data showed a good fit to the three dimensions structure of separate cognitive, emotional and somatic factors (*Chi^2^*=119; *df*=117; *p=*.4; CFI=.99; RMSEA=.006), reported earlier in the literature.

**Discussion:** Psychometric characteristics of the Dutch version of RPQ proved excellent to good, and can the instrument therefore be applied for research purposes and in daily clinical practice.

## Introduction

Traumatic brain injury (TBI) is one of the most common neurological conditions, and occurs when an external force causes brain trauma (1). TBI can be classified as mild, moderate or severe, and may have wide-ranging physical and psychological effects (2–7). Some signs or symptoms appear immediately after the traumatic event, while others days or weeks later. In the Netherlands, about 85,000 people are confronted with a traumatic brain injury, on a yearly basis (https://www.hersenstichting.nl/alles-over-hersenen/hersenaandoeningen/cijfers-over-patienten). On average 30,000 of these seek help at the Emergency Room (ER) of the hospital, and about 21,000 require hospital stays (8). Yearly, about 1,000 die because of TBI (9). Most people suffer from mild TBI (mTBI), e.g. concussion, for which they often do not seek professional help, or seek advice from their General Practitioner (GP): In the Netherlands, the GP functions as gatekeeper to the rest of the medical system. (8). Virtually all non-institutionalized Dutch citizens are registered with a GP controlling access to specialized medical care. (10)

In about one third of the cases mTBI leads to long-term consequences (5,11). A substantial proportion (about 15-30%) of individuals after mTBI are at risk to developing post-concussion symptoms (2). These can be classified into four categories: cognitive difficulties (e.g. concentration and memory loss), behavioral maladaptation (e.g. impulsivity, and aggressive behavior), psychiatric conditions (e.g. posttraumatic stress), and physical disorders (e.g. chronic pain) (11). Whether patients develop post-concussion symptoms, is associated with cognitive, emotional, behavioral and social risk factors, and does not necessarily depend on the severity of the traumatic brain injury, see fig 1 (2). Years of education, pre-injury psychiatric disorders, neck pain and prior TBI were found strong predictors of 6-month post-concussive symptoms (3,5), as were patient’s perceptions of their brain injury, their behavioral responses, passive and avoidant coping styles and emotional distress in response to this (2,5).

**Figure 1.**
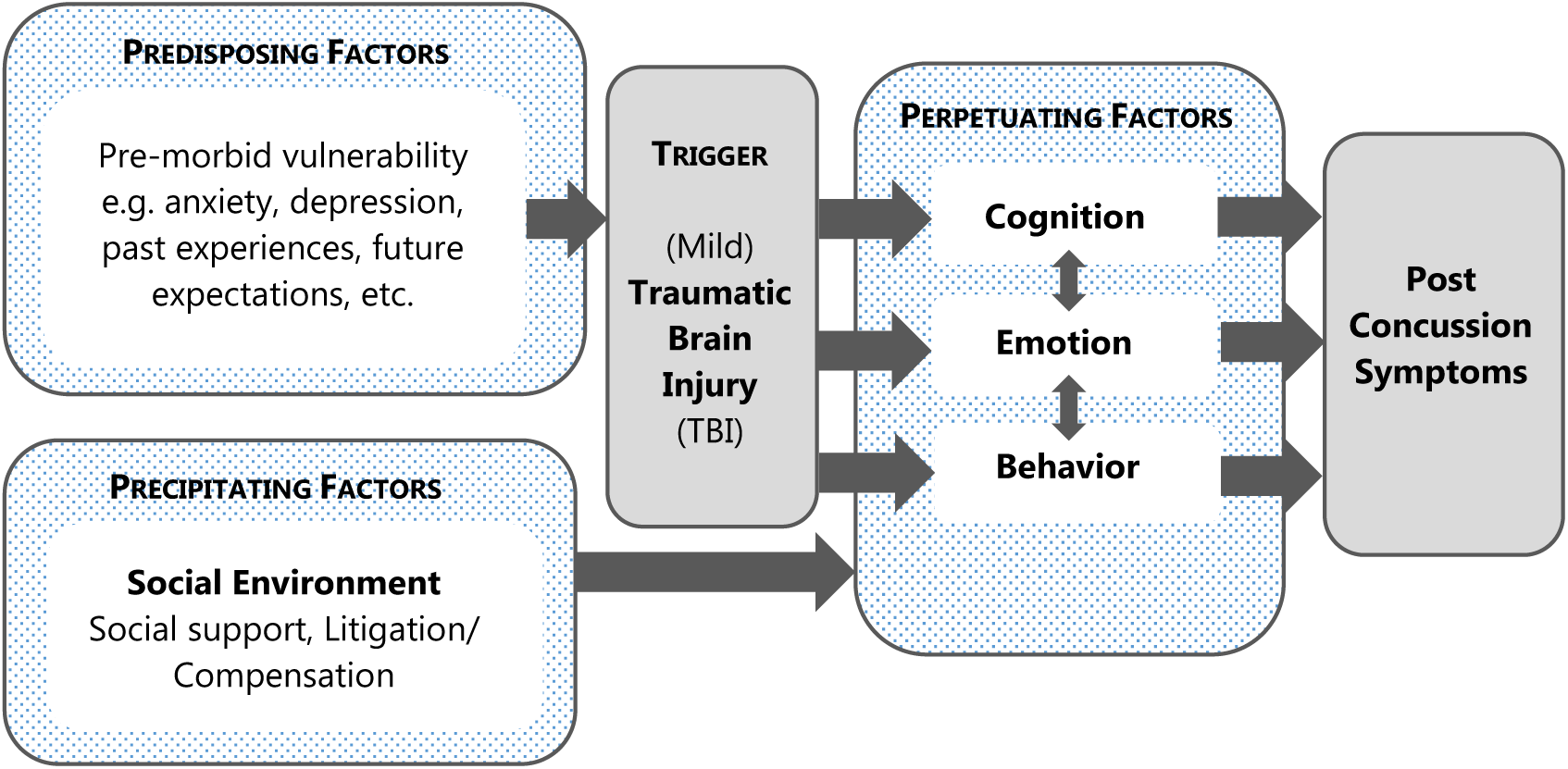
Factors influencing the development of Post-Concussion Symptoms after TBI (2)

As most people recover from their TBI within weeks to months after the injury, post-concussion symptoms might easily be overlooked, since these residual complaints may be deferred. Furthermore, imaging techniques often do not show any structural brain damage in this population (12,13). Still, one in three mTBI patients will not be able to resume work and activities six months after the event at a level similar to that before the accident(5). As a consequence, (m)TBI is associated with substantial ongoing disability and distress for patients, and high healthcare costs (2,8). A possible instrument for (early) identification in order to timely guide clinical management of post-concussion symptoms after TBI, is the Rivermead Post Concussion Syndrome Questionnaire (RPQ). The RPQ is a validated measurement-instrument to survey post-concussion symptoms, relying on self-report as to the presence and severity of 16 symptoms (14–17). The items form one scale, but were not always found to tap into the same underlying construct (3,4,14)). Eyres et al (2005) found no evidence for a single factor structure, and proposed to split the RPQ into two subscales consisting of the first three items ‘RPQ3’, representing immediate symptoms (headaches, dizziness, and nausea) and the remaining 13 items ‘RPQ13’, representing symptoms that might occur at a later stage. On the other hand, Lannsjö et al (2011) found strong support for both a single and two factor structure in their RPQ validation study, but failed to reproduce the RPQ3/13 two-category model as suggested by Eyres et al (2005). Furthermore, a ‘rationally-based’ three categories model was proposed by Smith-Seemiller and colleagues (2003), comprising of the following domains: 1. cognitive deficits (impaired memory, poor concentration, slow thinking), 2. somatic complaints (headaches, dizziness, nausea, blurred or double vision, noise or light sensitivity, sleep disturbance, fatigue), and 3. emotional complaints (irritably, depression, frustration, restlessness), serving as framework in various studies on post-concussion symptoms (3,18). The results of Potter and colleagues (2006) supported the existence of separate cognitive, emotional and somatic factors (17). So far, the RPQ has been validated in various languages (4,6), but until now this has not been the case for the Dutch version of this questionnaire. Therefore, this study aims to investigate the psychometric characteristics of the current Dutch translation of the RPQ.

## Methods

### Study sample

This study is part of the Collaborative European NeuroTrauma Effectiveness Research in TBI (CENTER-TBI) study, which is a prospective longitudinal observational study conducted in 72 centers from 21 countries (8). In the Netherlands, patients were recruited from seven medical centers spread over the country: Leiden University Medical Center (LUMC), University Medical Center Groningen (UMCG), Erasmus MC Rotterdam, Radboud University medical center Nijmegen, Medical center Haaglanden The Hague, Elisabeth Hospital Tilburg, HAGA hospitals The Hague. Furthermore, two centers in the Dutch-speaking-part of Belgium (Flanders) were included in the study: the Antwerp University Hospital, and the University Hospital in Leuven. Patients that presented within 24 hours after brain injury at the hospital, that were diagnosed with TBI, and had a clinical indication for CT scan, were eligible for the study, and were all invited to participate in this convenience sample. Those willing to participate provided written informed consent prior to inclusion. Patients with severe pre-existing neurological disorder that could confound the outcome assessment were excluded. A written informed consent to participate in the study was obtained at the time of inclusion. At six months post TBI, the nurse at the center administered the RPQ during a visit, or was sent by postal mail to those who did not need to attend the hospital, for completion at home. A pre-franked envelope was included to send it back.

### Translation of the Dutch RPQ

Two native Dutch speakers who are proficient in English translated the RPQ into Dutch, after which a native English speaker, who is fluent in Dutch, backward translated the harmonized version. This version was then compared to the original English RPQ version possible differences were identified and resolved by the two parties. In addition, a team of researchers and CENTER-TBI collaborators refined and reshaped the measurement-instrument until consensus was reached, following an iterative process. This multiple-step procedure resulted in a final version of the Dutch RPQ.

### Ethical Approval

The CENTER-TBI study (EC grant 602150) has been conducted in accordance with all relevant laws of the EU if directly applicable or of direct effect, and all relevant laws of the country where the Recruiting sites were located, including, but not limited to, the relevant privacy and data protection laws and regulations (the “Privacy Law”), the relevant laws and regulations on the use of human materials, and all relevant guidance relating to clinical studies from time to time in force including, but not limited to, the ICH Harmonised Tripartite Guideline for Good Clinical Practice (CPMP/ICH/135/95) (“ICH GCP”) and the World Medical Association Declaration of Helsinki entitled “Ethical Principles for Medical Research Involving Human Subjects”. Ethical approval was obtained for each recruiting site. Informed Consent was obtained for all patients recruited in the Core Dataset of CENTER-TBI and documented in the e-CRF. The list of sites, Ethical Committees, approval numbers and approval dates can be found on the official Center TBI website (www.center-tbi.eu/project/ethical-approval).

### Measurement-Instrument

The Rivermead Post-Concussion Questionnaire (RPQ) consists of 16 common symptoms related to post concussion. Patients are asked to rate how problematic symptoms were compared to the situation before the head injury on a 5-point Likert scale (0-4). A score of 0 indicating ‘not experienced at all; 1 indicating ‘no more of a problem (than before)’, 2 indicating ‘a mild problem’; 3 indicating ‘a moderate problem; 4 indicating ‘a severe problem’ (14). Scores are taken as sum of all symptom scores, excluding scores of 1, as these indicate symptoms are unchanged since the brain injury. This gives a potential total score range of 0 (representing no change in symptoms since the head injury) to 64 (most severe symptoms) (4). If more than 5 of the items were missing from the 16, a score was not calculated and treated as missing. The RPQ total score is calculated using prorating as imputation method, if up to one third of the items were missing. In addition, the RPQ scoring method of Stulemeijer et. al. (2008) was applied where a score of highest 2 (a mild problem) to at least 13 of the 16 items is defined a favorable outcome. Stulemeijer et al (2008) showed that 94% of non-brain-injured patients (wrist-, or ankle distortion) would meet this criterion (19).

Further, TBI severity was rated using the Glascow Coma Scale (GCS), with scores between 3-8 indicating severe, 9-12 moderate, and 13-15 mild TBI (20–24). The GCS was administered within the first 24 hours after the brain injury occurred. Current disability was assessed by administering the extended Glasgow Outcome Scale (GOSE) (25). GOSE scores were used to differentiate between patients with remaining severe disability (3–4), moderate disability (5–6), and good recovery (21,22). In addition, socio-demographic data were collected, including gender, age, working status, education level, etc.

### Analyses

The psychometric characteristics of the Dutch version of the RPQ were determined at six months post TBI, using SPSS version 24, AMOS version 24, and R version 3.3.3 to performing classical and modern test-theory analyses.

Internal consistency was determined by calculating Cronbach's alpha, including the scale if any item were deleted. To testing construct validity, Principal Axis Factoring (PAF) was done by unweighted least squares and oblimin rotation on the 16 RPQ items, exploring the underlying constructs in the Dutch version of the RPQ, as no consistent underlying factor structure has been established so far. Items were included if the factor loading was 0.5 or higher and if factor loadings on the other factors were 0.1 or lower. Confirmatory factor analysis (CFA) was used to examine the fit of the Dutch RPQ data to the various factor structures that had been described earlier in literature: For this, we used the single model factor, reflecting post-concussion symptoms as unitary entity (15); the RPQ3 and RPQ13 two factor model (4); and the three factor model (17,18).

Concurrent criterion validity was assessed by analyzing the influence of important covariates on RPQ scores (GCS, GOSE) using t-tests and one-way ANOVA. Descriptive analyses were performed for sociodemographic variables (gender, age, education level, etc.). Although people in The Netherlands and Flanders (Belgium) both speak Dutch, the language used differs, and words might have a different meaning. Therefore, all tests were performed both for the entire research sample and for each country separately where possible.

## Results

### Sample

In total 472 patients filled in the Dutch version of the RPQ at six months post TBI. Of these, 397 were administered in the Netherlands and 75 in Belgium. Twenty-five participants were aged under 18 (18 in the Netherlands, and seven in Belgium) and were excluded from this study. Country of residence was registered for 437 patients, who were either living in Belgium (N=67), or the Netherlands (N= 368), apart from two in Nepal. Not all respondents were born in the Netherlands or Belgium (see table 1), but since their understanding of the Dutch language was sufficient to fill in the RPQ, and since they currently were living in the Netherlands or Flanders, they were not excluded from the study. There were 277/ 447 (62%) male respondents (resp. 231/ 379 (60.9%) in the Netherlands, and 46/ 68 (67.6%) in Belgium). The vast majority of the study population (68.2%) belonged to the middle aged and older age groups (38% was aged 45-65, 30.2% 65 and up). More than half were higher educated (64.2%), and were either married or living together (56.6%). Nearly half were not, or no longer employed (48.4%), see table 2 for further details. There were no significant differences in total RPQ scores at six months post TBI for study participants in the Netherlands (*M*=12.63; *SD*=13.77) and participants in Belgium (*M*=12.64; *SD*=11.94; *t*(445)=-110*; p*=.9). The magnitude of the differences in the means was very small (eta squared<.0001).

**Table 1.** Overview of countries of birth of the respondents (N=447)

|  |  |  |  |  |  |
| --- | --- | --- | --- | --- | --- |
| Aruba<br>(N=1) | Brasil<br>(N=1) | Germany<br>(N=2) | Indonesia<br>(N=4) | Morocco<br>(N=3) | Surinam<br>(N=6) |
| Bosnia and<br>Herzegovina<br>(N=1) | China<br>(N=3) | Spain<br>(N=1) | Ireland<br>(N=1) | Netherlands<br>(N=326) | Saint-Martin<br>(N=1) |
| Belgium<br>(N=59) | Colombia<br>(N=1) | UK<br>(N=3) | Iran<br>(N=1) | Slovenia<br>(N=1) | Turkey<br>(N=3) |
| Bermuda<br>(N=1) | Cape Verde<br>(N=1) | Greece<br>(N=1) | Italy<br>(N=1) | Somalia<br>(N=1) | Vietnam<br>(N=1) |

**Table 2.** Patient Demographics, presenting percentages and numbers.

| **Agegroup** | **% NL (N)** | **% BE (N)** | **% Total (N)** |
| --- | --- | --- | --- |
| **18 to 24** | 12.9 (49) | 5.9 (4) | 11.9 (53) |
| **25 to 34** | 10.3 (39) | 11.8 (8) | 10.5 (47) |
| **35 to 44** | 8.2 (31) | 16.2 (11) | 9.4 (42) |
| **45 to 54** | 13.7 (52) | 19.1 (13) | 14.5 (65) |
| **55 to 64** | 23.2 (88) | 25.0 (17) | 23.5 (105) |
| **65 to 74** | 18.7 (71) | 13.2 (9) | 17.9 (80) |
| **75 and up** | 12.9 (49) | 8.8 (6) | 12.3 (55) |
| **Total (N)** | **379** | **68** | **447** |
| **Employment Status** | **% NL (N)** | **% BE (N)** | **% Total (N)** |
| **Working >34hpw** | 28.7 (118) | 7.3 (30) | 36.0 (148) |
| **Working 20-34hpw** | 9.2 (38) | 1.2 (5) | 10.5 (43) |
| **Working < 20hpw** | 3.6 (15) | .2 (1) | 3.9 (16) |
| **Currently sick leave** | 1.0 (4) | - | 1.0 (4) |
| **Special Employment** | .2 (1) | - | .2 (1) |
| **Unemployed** | 3.2 (13) | .7 (3) | 3.9 (16) |
| **Unable to work** | 2.2 (9) | .7 (3) | 2.9 (12) |
| **Retired** | 26.8 (110) | 4.9 (20) | 31.6 (130) |
| **Student** | 7.1 (29) | .7 (3) | 7.8 (32) |
| **Homemaker** |  |  | 2.2 (9) |
| **Total (N)** | **345** | **66** | **411** |
| *Missing (N)* | 34 | 2 | 36 |
| **Education Level** | **% NL (N)** | **% BE (N)** | **% Total (N)** |
| **None, not in school** | .3 (1) | .3 (1) | .5 (2) |
| **At school currently** | 2.8 (11) | - | 2.8 (11) |
| **Primary education** | 5.3 (21) | 1.5 (6) | 6.9 (27) |
| **Secondary education** | 17.3 (68) | 8.4 (33) | 25.7 (101) |
| **Post Highschool** | 33.1 (130) | 2.3 (9) | 35.4 (139) |
| **University/ College** | 24.7 (97) | 4.1 (16) | 28.8 (113) |
| **Total (N)** | **328** | **65** | **393** |
| *Missing (N)* | 51 | 3 | 54 |
| **Marital status** | **% NL (N)** | **% BE (N)** | **% Total (N)** |
| **Never Married** | 23.6 (99) | 3.6 (15) | 27.2 (114) |
| **Married** | 41.3 (173) | 7.9 (33) | 49.2 (206) |
| **Living Together** | 5.5 (23) | 1.9 (8) | 7.4 (31) |
| **Divorced/ Separated** | 8.6 (36) | 1.7 (7) | 10.3 (43) |
| **Widowed** | 5.0 (21) | .7 (3) | 5.7 (24) |
| **Total (N)** | **352** | **66** | **418** |
| *Missing (N)* | 27 | 2 | 49 |

At the time of the injury, 13.4% (N=60) of the study population solely attended the ER without further hospitalization. 52.6% (N=235) were hospitalized, and 34 % (N=152) needed a stay in the ICU. Initially, within 24 hours after TBI, 80.5% (N=316) were diagnosed mTBI, 5.8% (N=21) were diagnosed moderate TBI, and 6.4% (N=23) with severe TBI. Of 87 respondents (19.5%) these data were missing. Six months after the brain injury, GOSE scores reveal that 55.5% (N=248) of the respondents showed good recovery, 25% (N=112) reported moderate disabilities, and 6.7% (N=30) suffered from severe disabilities at that point in time. Of 57 respondents (12.8%) these data were missing. Those patients that solely attended the ER without further hospitalization, all were initially diagnosed mTBI (see Table 3a). Of 313 respondents all three data types were available (hospitalization, initial diagnosis and six months post recovery status), see table 3b for further descriptives.

**Table 3a.**
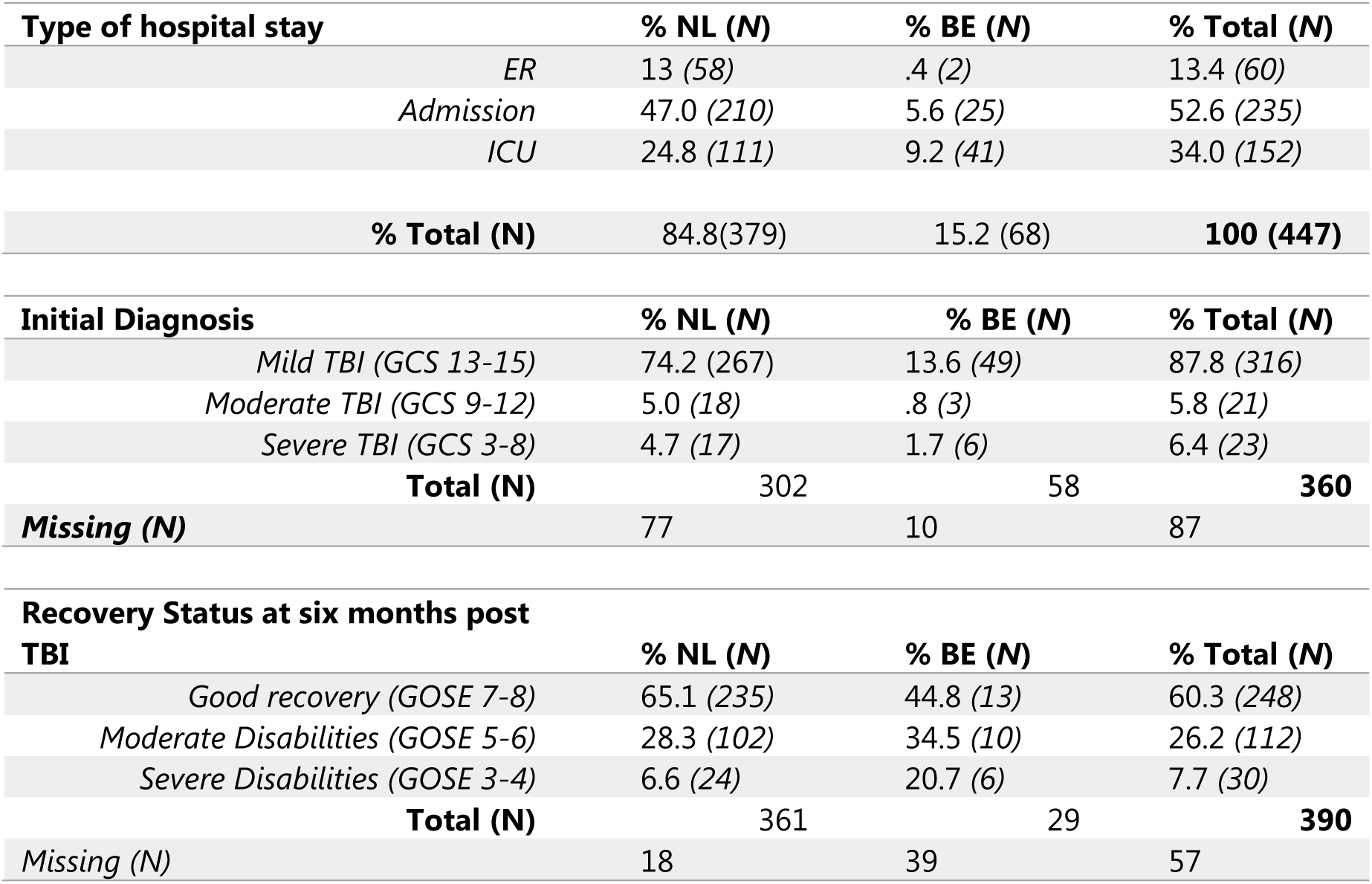
TBI-related patient demographics, presenting percentages and numbers.

| Type of hospital stay |  | % NL (N) | % BE (N) | % Total (N) |
| --- | --- | --- | --- | --- |
|  | ER | 13 (58) | .4 (2) | 13.4 (60) |
|  | Admission | 47.0 (210) | 5.6 (25) | 52.6 (235) |
|  | ICU | 24.8 (111) | 9.2 (41) | 34.0 (152) |
| % Total (N) |  | 84.8(379) | 15.2 (68) | 100 (447) |
| Initial Diagnosis |  | % NL (N) | % BE (N) | % Total (N) |
|  | Mild TBI (GCS 13-15) | 74.2 (267) | 13.6 (49) | 87.8 (316) |
|  | Moderate TBI (GCS 9-12) | 5.0 (18) | .8 (3) | 5.8 (21) |
|  | Severe TBI (GCS 3-8) | 4.7 (17) | 1.7 (6) | 6.4 (23) |
| Total (N) |  | 302 | 58 | 360 |
| Missing (N) |  | 77 | 10 | 87 |
| Recovery Status at six months post TBI |  | % NL (N) | % BE (N) | % Total (N) |
|  | Good recovery (GOSE 7-8) | 65.1 (235) | 44.8 (13) | 60.3 (248) |
|  | Moderate Disabilities (GOSE 5-6) | 28.3 (102) | 34.5 (10) | 26.2 (112) |
|  | Severe Disabilities (GOSE 3-4) | 6.6 (24) | 20.7 (6) | 7.7 (30) |
| Total (N) |  | 361 | 29 | 390 |
| Missing (N) |  | 18 | 39 | 57 |

**Table 3b.**
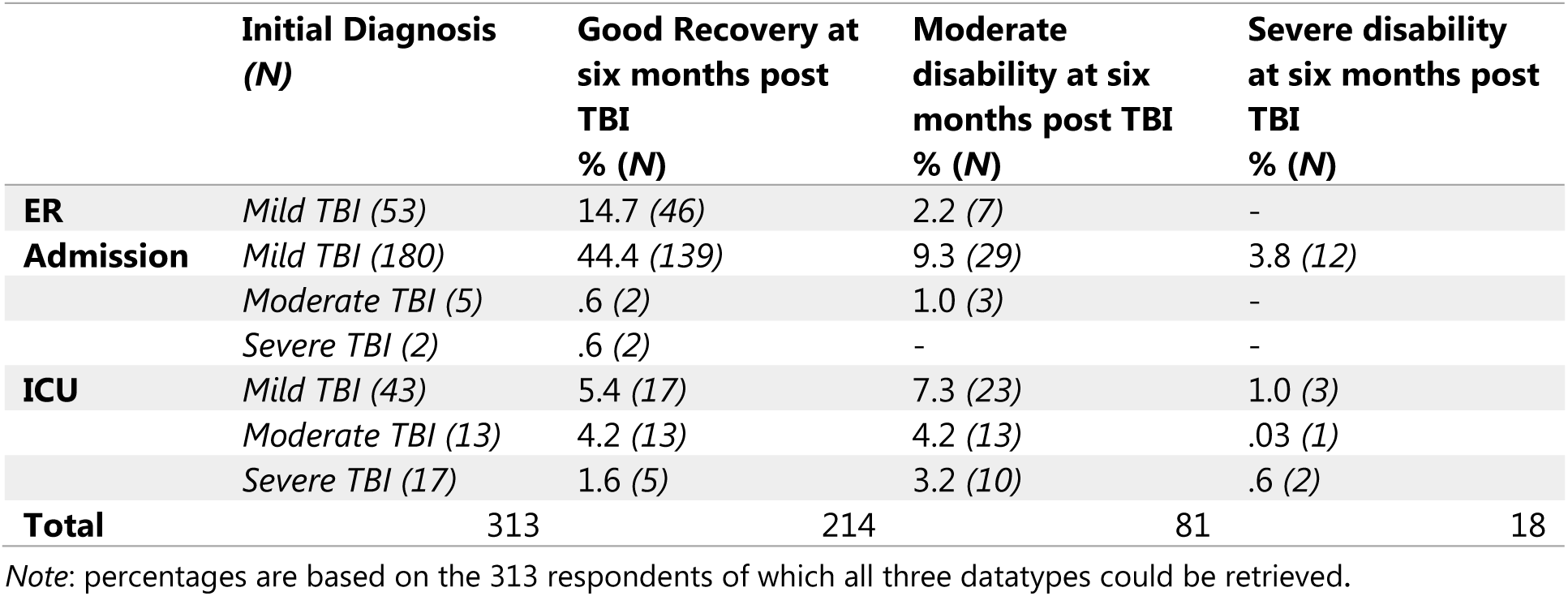
Patient recovery status at six months TBI by type of hospital stay and initial diagnosis.

### Factor analyses

Prior to performing PAF the suitability of data for factor analyses was assessed. Inspection of the correlation matrix revealed the presence of many coefficients of .3 and above. The Kaiser-Meyer-Oklin (KMO) value was .94, exceeding the recommended value of .6 (Kaiser 1970, 1974) and the Barlett’s Test of sphericity (Barlett, 1954) reached statistical significance (*p* <.0001), supporting the factorability of the correlation matrix. PAF, using Oblimin rotation, revealed the presence of three components with eigenvalue exceeding 1, explaining 47.7%, 5.0%, and 4.0% of the variance respectively. The scree plot revealed a clear break after the first component, see figure 2. Moreover, all items except for three (nausea (.42), blurred vision (.42) and double vision (.34)) show factor loadings of .5 and up on the first factor, but high factor loadings (>.1) on at least one of the other factors too. Confirmatory Factor analysis (CFA) was run to testing model fit to possible underlying factor structures that had been described in literature earlier (4,6,17,18). A central assumption is that the data are distributed normally. However, substantial problems with univariate skew and kurtosis were identified, see table 4. To correct for this data were dichotomized, computing 0 and 1 responses into 0, and 2, 3, and 4 into 1. Following this, CFA indicated a lack of fit to unitary model, given the significant Chi-squares (15) (*Chi^2^*=285.5; *df*=120; *p*<.001; CFI=.99; RMSEA=.06), and a lack of fit to the RPQ3/ RPQ13 two component model(4) (*Chi^2^*=271; *df*=119; *p*<.001; CFI=.99; RMSEA=.05), but showed a good fit to the three factor structure(18) (*Chi^2^*=119; *df*=117; *p=*.4; CFI=.99; RMSEA=.006). The Belgian sample was too small to performing separate CFAs.

**Table 4.**
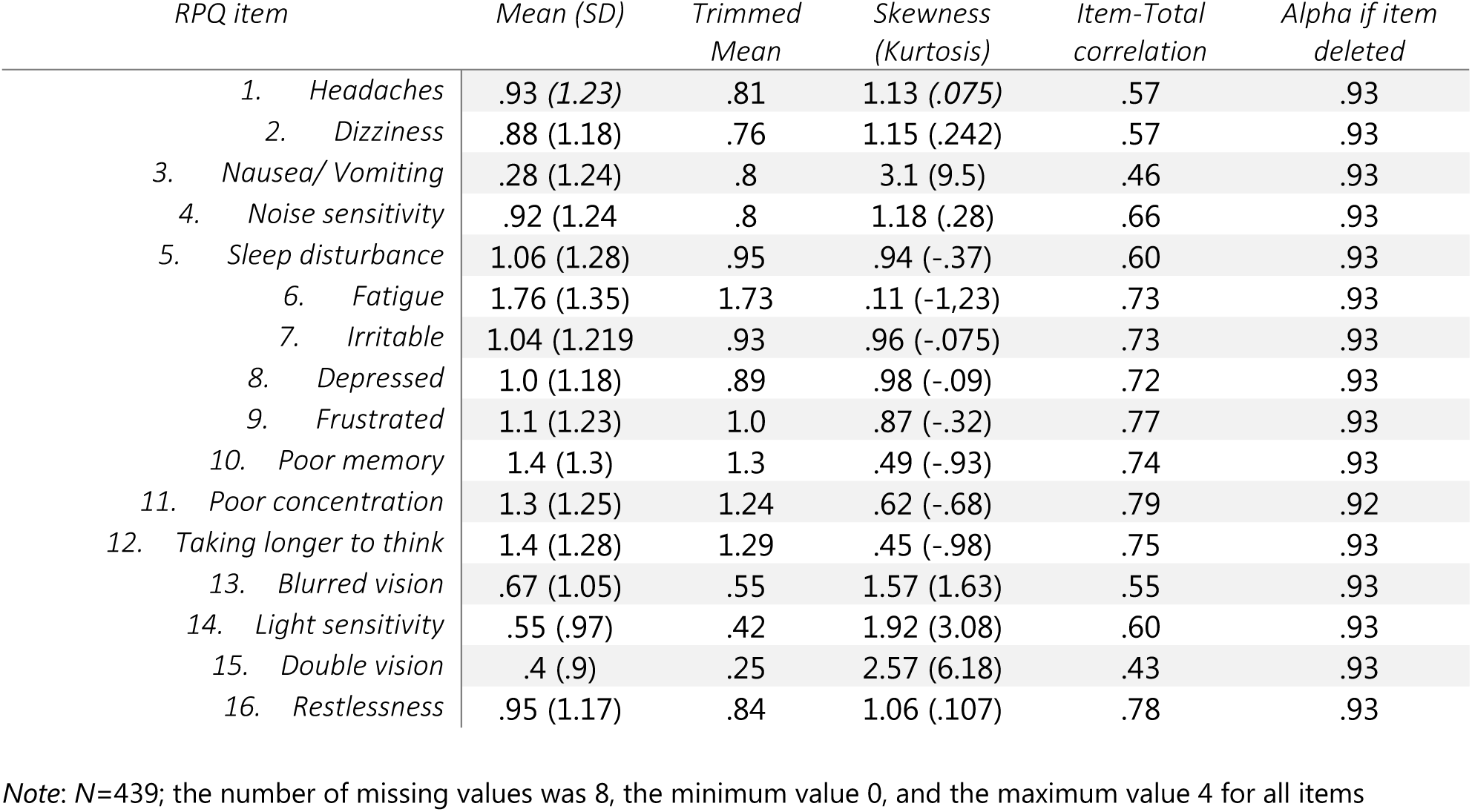
individual item descriptives and characteristics.

| *RPQ item* | *Mean (SD)* | *Trimmed Mean* | *Skewness (Kurtosis)* | *Item-Total correlation* | *Alpha if item deleted* |
| --- | --- | --- | --- | --- | --- |
| 1. Headaches | .93 (1.23) | .81 | 1.13 (.075) | .57 | .93 |
| 2. Dizziness | .88 (1.18) | .76 | 1.15 (.242) | .57 | .93 |
| 3. Nausea/ Vomiting | .28 (1.24) | .8 | 3.1 (9.5) | .46 | .93 |
| 4. Noise sensitivity | .92 (1.24) | .8 | 1.18 (.28) | .66 | .93 |
| 5. Sleep disturbance | 1.06 (1.28) | .95 | .94 (-.37) | .60 | .93 |
| 6. Fatigue | 1.76 (1.35) | 1.73 | .11 (-1.23) | .73 | .93 |
| 7. Irritable | 1.04 (1.219) | .93 | .96 (-.075) | .73 | .93 |
| 8. Depressed | 1.0 (1.18) | .89 | .98 (-.09) | .72 | .93 |
| 9. Frustrated | 1.1 (1.23) | 1.0 | .87 (-.32) | .77 | .93 |
| 10. Poor memory | 1.4 (1.3) | 1.3 | .49 (-.93) | .74 | .93 |
| 11. Poor concentration | 1.3 (1.25) | 1.24 | .62 (-.68) | .79 | .92 |
| 12. Taking longer to think | 1.4 (1.28) | 1.29 | .45 (-.98) | .75 | .93 |
| 13. Blurred vision | .67 (1.05) | .55 | 1.57 (1.63) | .55 | .93 |
| 14. Light sensitivity | .55 (.97) | .42 | 1.92 (3.08) | .60 | .93 |
| 15. Double vision | .4 (.9) | .25 | 2.57 (6.18) | .43 | .93 |
| 16. Restlessness | .95 (1.17) | .84 | 1.06 (.107) | .78 | .93 |

Note: N=439; the number of missing values was 8, the minimum value 0, and the maximum value 4 for all items

**Figure 2.**
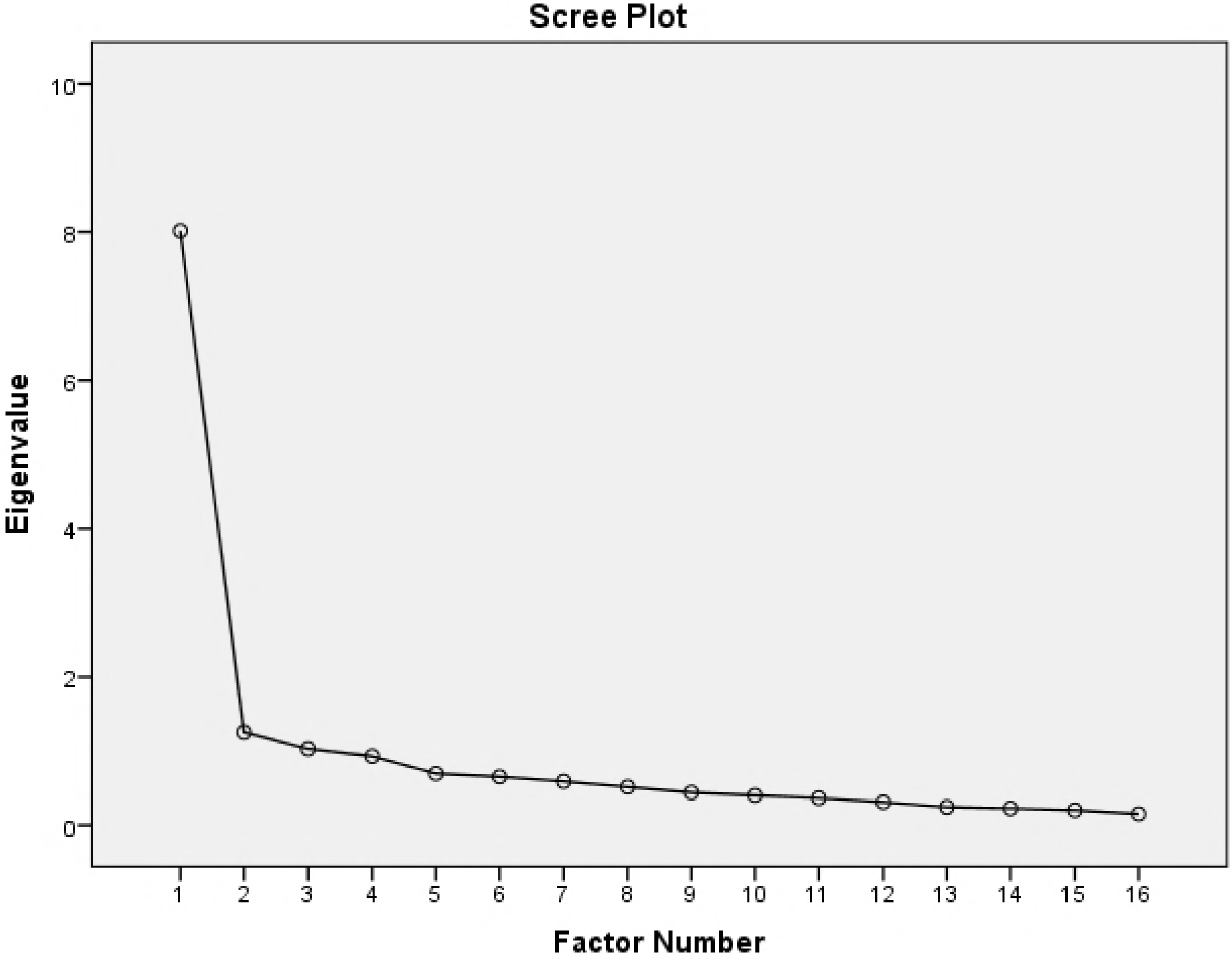
Visual representation of factor loadings

### Quality Criteria

The RPQ showed high consistency with Cronbach’s alpha being .93. For the sample in the Netherlands Cronbach’s alpha was .94, and for the sample in Belgium Cronbach’s was alpha .91. The scale did not improve if any items were deleted. Spearman Brown Coefficient *r_sb1 was_* .91). Further, item characteristics showed high item correlations (>.55, except for double vision and nausea), and acceptable asymmetry for all items but double vision (skewness being 2.6), see table 5

### Concurrent Criterion validity

RPQ total scores at six months post TBI of patients (self-)reporting severe (*M*=20.7; *SD*=18.3; *N*=30) or moderate (*M*=20.2; *SD*=13.9; *N*=112) disabilities according to their total scores on the GOSE scale, differed significantly to those that showed good recovery at this point in time (*M*=8.3; *SD*=10.6; *N*=248) (*F(2, 387)*=42.7; *p*<.001). RPQ total scores at six months post TBI further were found to differentiate between initial mTBI (*M*=11.6; *SD*=13.2; *N*=316) and moderate TBI (*M*=20.2; *SD*=16.8; *N*=21) diagnoses (GCS-scores) (*F(2, 357)*=4.5; *p*=.012). Remarkably, RPQ total scores of those initially diagnosed with severe TBI (*M*=14.8, SD =13.9; *N*=23), resembled most those initially diagnosed with mTBI (*NS*). When recalculating RPQ total score into favorable (a score of highest 2 to at least 13 of the 16 items (26)) vs unfavorable, 74.7% (*N*=334) of the study population had a favorable outcome at six months post TBI, indicating that the symptoms reported, do not differ from what can be found in a non-TBI population (26,27). 25.3% (*N*=113) still had an unfavorable, strongly related to TBI, outcome. The RPQ score was found to discriminate between recovery status (GOSE scores) at six months post TBI (*Chi^2^*=45.2; *df*=2; p<.001), although not strongly (*Cramer’s V* =.11). When solely taking the sample from the Netherlands into account, a stronger relationship between favorable and unfavorable RPQ outcomes and recovery status (GOSE scores) at six month post TBI was found (*Cramer’s V* =.35, *Chi^2^*=43.8; *df*=2; p<.001). Further, the RPQ total scores at six months post TBI of the Dutch sample were found to discriminate between recovery status (GOSE scores) at six months post TBI (*F(2, 358)*=39.2; *p*<.001), and initial diagnoses within 24 hours after the brain injury occurred (GCS Scores) *(F(2, 299)*=3.7; *p*=.3). The number of participants from Belgium that could be included in these analyses were too low for further analyses.

## Discussion

The current Dutch translationn of the RPQ showed good psychometric characteristics, with high internal consistency, and good construct validity. As for these aspects, it can be applied for research purposes and in daily clinical practice, as an instrument to identify post-concussion symptoms. Besides, it proved sensitive for recovery status at six months post TBI, showing that those who (self-) reported moderate or severe disabilities six months after the brain injury took place, had significant lower RPQ total scores compared to those reporting good recovery at that time point. Further, RPQ total scores at six months post TBI were found to distinguish between initial TBI diagnoses: Those initially diagnosed with moderate TBI had higher RPQ total (sum) scores at six months post TBI compared to those initially diagnosed with mild TBI (mTBI). However, the number of people in our research sample that were initially diagnosed with moderate TBI was low. Moreover, RPQ total scores at six months post TBI of those initially diagnosed with severe TBI resembled more the total score of those diagnosed with mTBI, rather than those with moderate TBI. A possible explanation for this might be that moderate TBI and the amount of care needed was underestimated. This type of TBI might need more intensive care than what was provided. However, again the number of people in this group was too low to base further conclusions upon. Another explanation for this might be that the large number of mTBI-diagnosed patients who were admitted to the ICU, needed intensive care because of other injuries, and thus were diagnosed mTBI correctly.

Consistent with the findings of others, we found multidimensionality as underlying structure of the RPQ measurement-instrument (3,4,6,17,18,26). However, high factor loadings of items on multiple factors, and the clear break after the first factor in the scree plot, would suggest a one factor structure rather than multidimensionality. Confirmative factor analyses on the other hand revealed that our data would fit best to the three-component model dividing the RPQ post-concussion symptoms into the following three categories: 1. cognitive deficits (impaired memory, poor concentration, slow thinking), 2. somatic complaints (headaches, dizziness, nausea, blurred or double vision, noise or light sensitivity, sleep disturbances), and 3. emotional complaints (irritably, depression, frustration, restlessness). Small differences were found between the Belgium and Dutch sample, with only the Dutch sample showing a good fit to this model. However the number of respondents in the Belgium sample was below 250, due to which the criteria for model fit may not be valid (28).

The variation in underlying structure of the RPQ differs between studies and countries and might be due to various reasons. One reason could be the convenience sample used for this research. Another explanation might be the different analyses techniques used in the various studies, as modern techniques often tend to disqualify measurement-instrument validity that had been established before by classical analyses methods (29). A third possible explanation underpinning this may be the way in which measurement-instruments are being translated in accordance to the WHO guidelines of forward and back translation (30). In order not to lose the potential to comparing data, researchers prefer to stay as close as possible to the original version. However, through this, the principles of cultural interpretation and translating the correct meaning of the items, might be missed out, due to which country differences might occur, even though the instrument used is very similar. (31,32).

In addition, we argue that despite the underlying multidimensionality found, the Dutch version of the RPQ needs not necessarily be divided into subscales when applied for research purposes and in daily clinical practice. The underlying multidimensionality might indicate that post-concussion symptoms represent more than one dimension, but factors highly correlated, and items were not unique for just one of the factors. Moreover, there is a large body of evidence that supports the use of total scores of scales to which multidimensionality is a precondition, e.g. attitude scales that usually exist of a cognitive and affective component(33). Furthermore, as the psychometric properties of the Dutch version of the RPQ proved good, it would be of interest to implement this measurement-instrument into primary care settings, in order to timely recognize the possible long-term consequences of TBI. This would especially be effective in countries as the Netherlands where one has to see the GP first, before entering the rest of the medical system, the so-called gatekeeper system, and in ER settings, where people usually are only checked medically and then send home, in order to timely identify potential patients at risk. However, more clarity is needed on how to best interpret RPQ scores(17), since similar symptoms can too be reported by those suffering from different injuries and, disorders, or by members of the general population as fatigue, headaches, nausea etc., are very common. As such, Stulemeijer and colleague (2016) found that 94% of non-TBI patients with wrist or ankle distortion too score positive on a maximum of three RPQ items.

### Limitations

At six months post TBI, three quarters of the research sample no longer showed post-concussion symptoms, due to which data were skewed and not normally distributed. Validation of the RPQ at this point in time might therefore be difficult. Further, items were strongly correlated, due to which items strongly loaded on one and the same factor. Another limitation of this study was the limited Belgian sample, which was often too small for sound complex analyses, such as CFA. Further, the lacking of a construct validation phase making use of cognitive interviewing limits the overall conclusion on validity of the Dutch version of the RPQ, especially since there were between-country differences. Knowing what our respondents think we are asking, and knowing how they interpret the questions we are asking might help to explain the variety in underlying symptom structure found too (29). Moreover, the response scale used (0-4) might be confusing since patients might find the following order of the score of zero indicating ‘no problem’, and the score of one indicating ‘no more of a problem’ difficult to understand, as a more natural following order would be: zero indicating no problem, and one indicating a small problem. Last, the convenience hospital sample used in this study might be limited representative to the entire mTBI population, as most people in the Netherlands tend not to seek specialized medical help for their head injury.

## Conclusion

The psychometric characteristics of the Dutch version of RPQ proved good, showing high consistency, and good construct validity, being sensitive to self-reported recovery status at six months post TBI and initial TBI-diagnosis sensitive. The Dutch version of the RPQ can therefore be applied for research purposes and in daily clinical practice. Further discussion is needed with regard to the scoring of the RPQ, as underlying multidimensionality may not necessarily stand in the way of using a total score.

## Acknowledgements

The research leading to these results was supported by the European Union’s Seventh Framework Programme (*FP7/2007-2013*) under *grant agreement* n° 602150 (CENTER-TBI). Additional support for CENTER-TBI was received from the Hannelore Kohl Stiftung. The authors would like to thank all patients that were willing to participate in the study and all CENTER-TBI investigators and their staff for completing the provider profiling questionnaires. Those particularly involved in this paper are listed underneath.

## Data Availability

There are however legal constraints that prohibit us from making the data available. Since there are only a limited number of centers per country included in this study (for two countries only one center), data will be identifiable. Readers may contact Dr Hester Lingsma for requests for the data.

